# Decoding Arbitrary and Informed Decisions from Intracranial Recordings in Humans

**DOI:** 10.1101/2023.06.01.543070

**Authors:** Laura Marras, Maxime Verwoert, Maarten C. Ottenhoff, Sophocles Goulis, Johannes P. van Dijk, Simon Tousseyn, Louis Wagner, Albert J. Colon, Pieter L. Kubben, Marcus L.F. Janssen, Steffen A. Herff, Christian Herff

## Abstract

Ideally, decisions are made based on prior knowledge, which allows for informed choices. Real life, however, often requires us to make decisions arbitrarily, without sufficient information. Decoding decision making processes from neural activity could allow for cognitive neuroprostheses and Brain-Computer Interfaces (BCIs) to support decision processes in rapid human-machine interactions, weigh decision-making confidence, and further enable neuromodulation protocols for the treatment of reward-related dysfunctions. To understand the differences between the decision-making processes in arbitrary and informed decisions, we recorded intracranial electroencephalography in a large number of cortical and subcortical areas from 5 patients during a categorization task. We demonstrate that individual decisions can be decoded from Local Field Potentials (LFPs) before motor response, in both arbitrary and informed conditions. Our analysis revealed dissimilar spatio-temporal patterns between arbitrary and informed decision-making, with arbitrary decisions being decodable in fewer brain regions and earlier in time compared to informed decisions.

## Introduction

Adaptive behaviors, intended as actions aimed at maximizing survival and minimizing harm, require the ability to evaluate environmental stimuli, process the possible responses, anticipate their consequences, and choose the most appropriate action (***Rangel et al., 2008***). Therefore, the capacity to make informed decisions is crucial for survival. Unfortunately, daily life is commonly characterized by novel situations where prior knowledge of how possible choices are mapped onto outcomes is not available. In such situations, initial reflexes and habits, followed by subsequent instrumental learning play a crucial role in creating action-outcome mappings through reward-driven exploration of the decision space (***Balleine, 2019***; ***O’Doherty et al., 2017***; ***Morris et al., 2022***). In this trial-and-error strategy, decisions are initially arbitrary, but the information obtained through the initial decisions can later be used to guide informed decision-making.

The neural underpinnings of informed and arbitrary decision-making remain uncertain. Studies that have examined these forms of decision-making individually suggest they may recruit distinct neural circuits and mechanisms (***Mudrik et al., 2019***). In particular, informed decisions have been associated with activity in the prefrontal cortex (***Wallis and Miller, 2003***), orbitofrontal cortex (***Wallis et al., 2007***), and anterior cingulate cortex (***Kennerley et al., 2006***; ***Hart et al., 2020***), whereas arbitrary decisions have been linked to activations in the supplementary motor area, medial frontal cortex and parietal cortex (***Soon et al., 2008***; ***Fried et al., 2011***; ***Soon et al., 2013***; ***Tosoni et al., 2008***). However, conflicting evidence emerges from studies that have directly compared the electrophysiological responses associated with these distinct decision-making processes (***Bold et al., 2022***). Some studies found that the Readiness Potential (RP) - a negative electrical component associated with motor decisions (***Schurger et al., 2021***) - is smaller or absent in informed compared to arbitrary decisions (***Maoz et al., 2019***; ***Travers and Haggard, 2021***). In contrast, other studies reported the opposite effect (***Travers et al., 2021***) or no differences at all (***Parés-Pujolràs et al., 2021***).

Understanding the neurophysiological signatures of informed and arbitrary decisions represents the first step toward the development of neural prosthetics such as targeted neuromodulation protocols (***Orsini et al., 2019***) and decision-making Brain-Computer Interfaces (BCIs) (***Mirabella and Lebedev, 2017***; ***Andersen et al., 2010***; ***Pesaran et al., 2006***). Specifically, closed-loop neuromodulation techniques have been revealed to be promising in the treatment of disorders characterized by deficits in reward-related behaviors and decision-making, such as addiction (***Habelt et al., 2020***; ***Luigjes et al., 2019***; ***Fecteau et al., 2010***), pathological gambling (***Fecteau et al., 2010***; ***Goudriaan and Schluter, 2019***), and affective disorders (***Bina and Langevin, 2018***). However, these approaches require the identification of reliable neural features of decision-making that can be robustly decoded and are suitable to serve as control signals for stimulation. Additionally, being able to decode decisions early, before motor planning onset, might contribute to the improvement of BCIs for motor control and communication, allowing shorter reaction times and better error processing (***Mirabella and Lebedev, 2017***).

Previous studies in Non-Human Primates have shown that decision-related features can be decoded from electrophysiological recordings. Rich and Wallis were able to decode option values during binary decisions, based on single-unit activity (SUA) and local field potential (LFP) recordings from the monkey orbitofrontal cortex (***Rich and Wallis, 2016***). In a recent study, a linear decoder was applied to neural population activity from the primary motor and dorsal premotor cortices to track the deliberation process and predict the animal’s choices at the single trial level (***Peixoto et al., 2021***). These studies demonstrated the feasibility of single-trial characterization of neural features in decision-making based on small- and large-scale electric signals in animals.

In humans, early attempts at decoding decision-related variables were carried out in functional magnetic resonance imaging (fMRI) studies. Specifically, by applying multivariate techniques to fMRI recordings, multiple studies achieved successful prediction of various behavioral aspects of decision-making, including reward-related decisions (***Hampton and O’doherty, 2007***), belief or disbelief decisions (***Douglas et al., 2013***), action intentions (***Gallivan et al., 2011***), and single-choice outcomes (***Haynes, 2011***). However, while fMRI allows the identification of spatial activation patterns underlying decision encoding, it does not reveal insights into the temporal dynamics or the role of oscillatory activity. Furthermore, the costs, lack of portability, and low temporal resolution render fMRI unfeasible for online BCIs or treatment solutions.

To this end, various studies have explored decoding single decisions from electroencephalography (EEG) activity in humans. In a study by Si and colleagues, the authors developed a computational framework to extract features from single-trial EEG activity and use them to predict individual decision responses in the ultimatum game (***Si et al., 2020***). They demonstrated that different decision responses could be decoded from EEG network activity on a trial-by-trial basis, supporting the idea that acceptance and rejection decisions engage different brain networks (***Si et al., 2019***). Similarly, a recent study implemented a novel feature extraction method to discriminate cooperation and aggression decisions based on EEG. The authors showed that this method can accurately classify different decision responses and that the primary source of discriminability comes from the EEG information within the frequency range below 40 Hz (***Huang et al., 2022***). In a different study, the authors were able to decode the intention to move left or right index based on EEG patterns extracted from a combination of movement execution, imagery, and preparation (***Salvaris and Haggard, 2014***). These findings support the role of specific rhythmic activity in decision encoding and suggest that neural fingerprints of this process can be extracted from electrophysiological activity. However, scalp EEG suffers from multiple limitations related to spatial resolution, signal-to-noise ratio, sensitivity to artifacts, and, most importantly, access to higher frequency bands (***Herff et al., 2020***; ***Buzsáki et al., 2012***; ***Parvizi and Kastner, 2018***), which substantially restrain its use as a potential signal for BCIs and neuroprosthetics.

Conversely, stereotactic EEG (sEEG) provides access to high-frequency LFP activity and allows to characterize the temporal patterns of cognitive processing within spatially well-defined brain sites (***Parvizi and Kastner, 2018***; ***Herff et al., 2020***). In particular, high-frequency LFP power has been indicated as an optimal signal for BCI control (***Zhuang et al., 2010***; ***Brincat et al., 2013***), as it reflects integrated spiking of multiple neurons near the recording site (***Buzsáki et al., 2012***; ***Ray et al., 2008***; ***Ray and Maunsell, 2011***) and has been shown to have comparable decoding accuracy and informative content with single-unit spikes (***Mehring et al., 2003***; ***Brincat et al., 2013***). Additionally, compared to SUA, using LFPs allows for up-scaling to more complex decision spaces whilst remaining in a timescale useful for BCIs, requires less computational power, and a lower data sampling rate (***Brincat et al., 2013***). Thiery and colleagues recorded sEEG while patients performed an oculomotor decision task involving free and instructed eye movements (***Thiery et al., 2020***). They showed that LFPs could be used to accurately classify saccades as instructed or free on a single-trial basis. Notably, while the brain areas engaged by free-choice and instructed actions were mostly shared, the two conditions differed in the temporal dynamics of high-frequency activity, which was found to be the most informative feature for the classification task.

Taken together, human and animal studies provide converging evidence that neural signatures of decision-making can be identified in broadband electrophysiological activity, which is best suited to extract reliable features for BCI control and neuroprosthetics applications.

In this study, we probed neurophysiological fingerprints of decision-making, focusing for the first time on informed and arbitrary decisions. Stereotactic EEG was recorded from a total of 590 contacts in 5 patients suffering from pharmaco-resistant epilepsy, during a decision-making paradigm including arbitrary and informed choices. By applying decoding analysis to broadband LFP activity, we show that the outcome of single categorization choices can be decoded with high accuracy for both arbitrary and informed decisions. Furthermore, the resolution of intracranial recordings allowed us to examine the spatial and temporal dynamics of decision decoding and identify previously unknown differences between informed and arbitrary decision-making.

## Results

Participants performed a simple feedback-guided categorization task in which they had to assign each presented stimulus to one of two abstract categories (i.e., “Winning” or “Losing”) using a trial-and-error strategy. Each stimulus was presented three times in a pseudo-randomized order (Fig.1a). Following the stimulus, participants were given 2 seconds to indicate the selected category by pressing the corresponding button (left or right arrow). Participants received feedback after each decision indicating whether they selected the correct category (Fig.1b). Since stimulus-category associations were random and meaningless, no task-relevant prior knowledge was available to the participants when performing the first trial of each stimulus. Therefore, whenever a new stimulus was presented, they had to simply guess (arbitrary decisions). Conversely, when stimuli were presented for the second and third time, participants were instructed to adjust their decisions based on the previously received feedback (informed decisions). Accordingly, we expected participants’ performance to be at chance level during arbitrary decisions and then improve during informed decisions.

**Figure 1.**
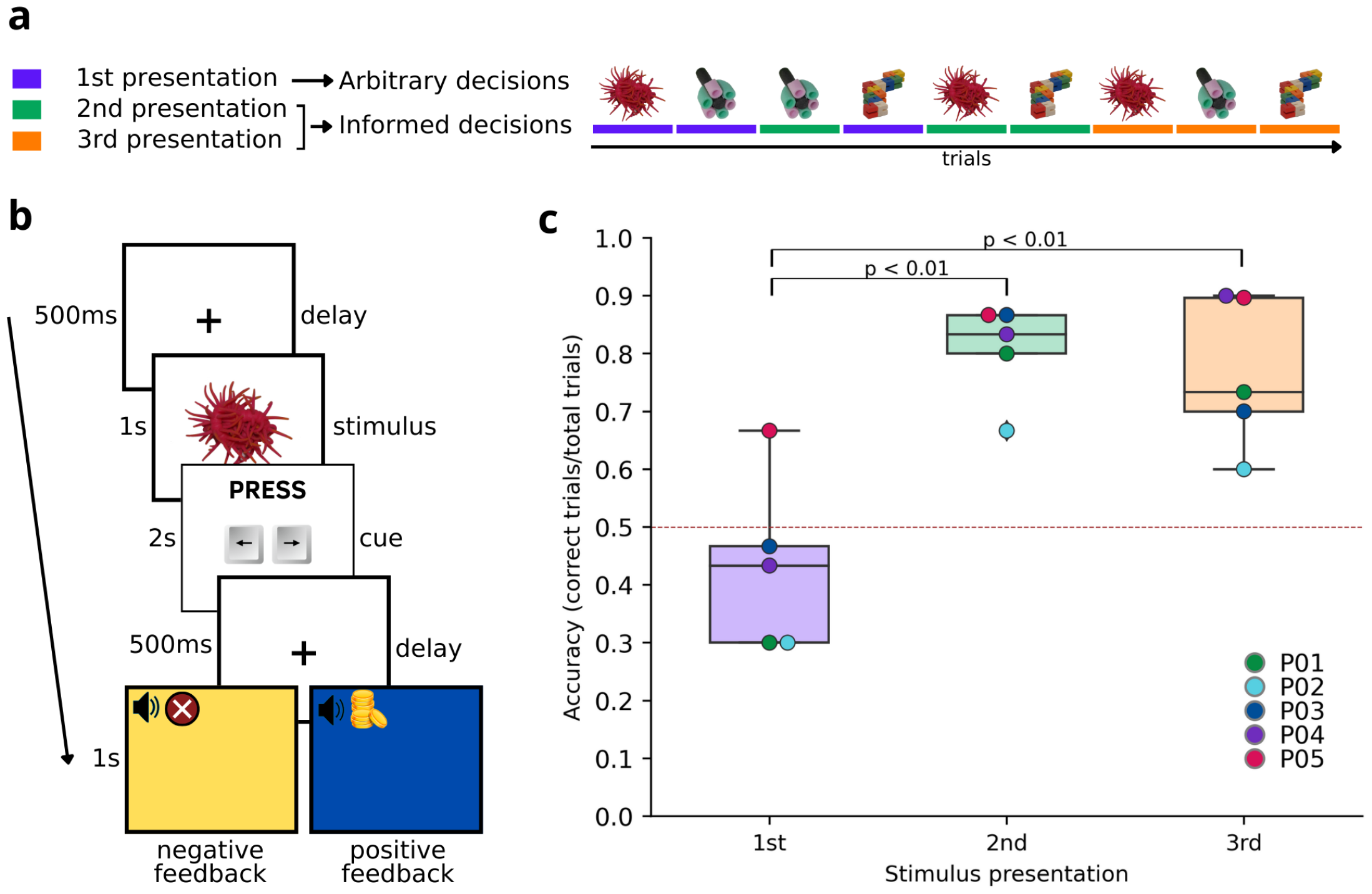
Feedback-guided categorization task. **a, *Schematic of an example session***. In each session, three visual stimuli are presented three times, in randomized order. Color indicates first (purple), second (green), and third (orange) stimulus presentations. Arbitrary decision condition consists of trials in which stimuli were presented for the first time and participants were instructed to guess their category. Informed decision condition includes trials with second and third presentations of stimuli, where participants had to adjust or confirm their choices based on feedback received in previous trials. **b, *Schematic of example trial sequence***. After 500ms of fixation period, a stimulus is presented for one second. Then a cue indicating to press the left (to choose “Winning” category) or right (to choose “Losing” category) arrow key appears for maximum 2s. As soon as a response is provided, a fixation cross is displayed for 500ms. Positive (blue screen and coins dropping sound) or negative (yellow screen and buzzing sound) feedback is delivered depending on whether participants chose the correct stimulus-category or not, presented for 1s. **c, *Task performance per participant and stimulus presentation***. Boxplots represent performance distribution for each stimulus presentation, each dot represents one participant’s average performance. The red dotted line indicates chance level.

### Participants shift from arbitrary to informed decisions in a categorization task

First, we tested whether feedback information was processed and integrated into the decisionmaking process. As expected, average accuracy across participants was at chance level (0.43) during arbitrary decisions. Performance significantly improved in the second (0.80, *p* < 0.01, Tukey’s HSD test) and third (0.76, *p* < 0.01, Tukey’s HSD test) compared to the first stimulus presentation (Fig.1c). These results show that participants successfully learned the correct stimulus-category association after feedback presentation and used the provided feedback information to improve their decisions. Average reaction time across participants was 335ms ±228ms after cue onset (428ms ±279ms in 1st presentation, 244ms ±188ms in 2nd presentation, and 333ms ±229ms in 3rd presentation).

### Decoding arbitrary and informed decisions from LFP activity

Secondly, we explored whether individual upcoming decisions could be decoded from single-channel LFP activity. We trained a linear discriminant analysis (LDA) classifier on multi-band LFP power averaged over the time period of stimulus presentation (Methods). We tested decoding of decision (selected category) for each recording channel of each participant and decision condition (arbitrary or informed). Permutation test revealed that out of a total of 590 channels, the classifier was able to decode individual choices with significant accuracy (ROC AUC, *p* < 0.05, permutation tests, T-Max correction) in 35 channels during informed decisions and 16 channels in the arbitrary decision condition (Fig.2a). Among these, performances varied across participants and channels, with ROC AUC ranging from 0.62 to 0.77. Notably, significantly accurate decision decoding was observed in at least one channel in every single participant, in both arbitrary and informed conditions. These results show that decoding decisions from multi-band LFP activity is feasible at the single channel level. This means that an upcoming decision can be predicted solely based on neural activity.

**Figure 2.**
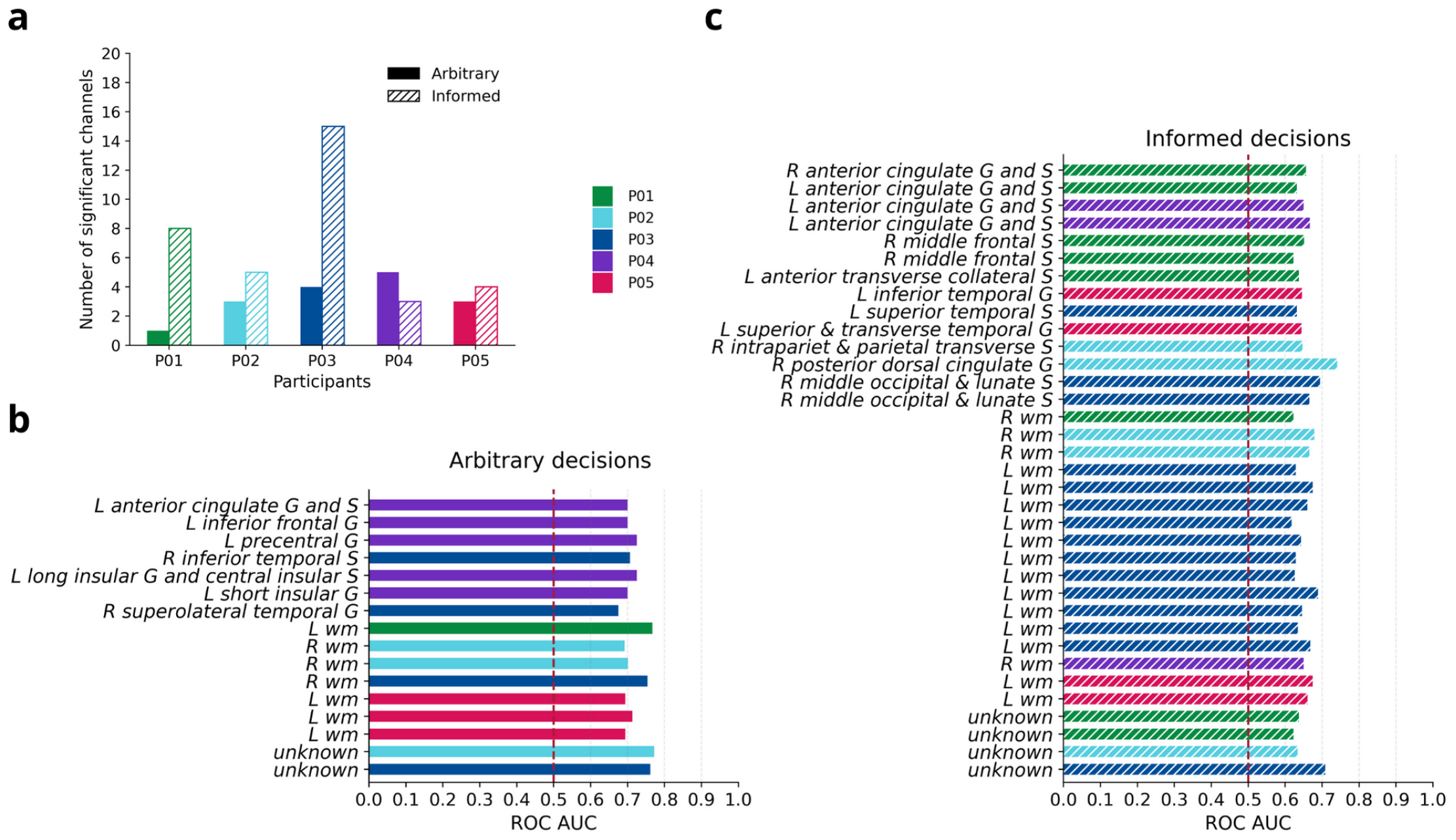
Decoding single-choice from single-channel multi-band LFP activity. An LDA classifier was applied to the multi-band LFP power (average over the entire stimulus period) of single channels to decode single decisions (“L” vs “W” presses), for each participant and condition separately. **a, *Number of informative channels per participant and condition***. For each participant (color coded) the number of channels that resulted in significant decoding ROC AUC (*p* < 0.05, permutation tests, T-Max correction) is shown for arbitrary (full bars) and informed (dashed bars) conditions. **b, *Decoding results of informative channels during arbitrary decisions***. The bar plot shows ROC AUC scores and anatomical location of all channels that resulted in significant decoding results (*p* < 0.05, permutation tests, T-Max correction) during arbitrary decisions. Bar colors represent participants as in **a**. L=left, R=right, G=gyrus, S=sulcus, wm=white matter. **c**, Same as in **b**. for the informed decision condition.

### Spatial dynamics of decision decoding

We then looked at the anatomical location of the contacts with significant decoding accuracy and compared the results across the two decision conditions. Across all participants, only 1 channel, located in the left anterior cingulate cortex, showed significant decoding performance in both conditions (Fig.2b, Fig.3). The 15 contacts that showed significant activity only during arbitrary decisionmaking were located in various portions of temporal, insular, and frontal cortices and in white matter (Fig.2b, Fig.3). Finally, the 34 channels that reached significant decoding performance exclusively for informed decisions were spread out across all four cortical lobes, including anterior and posterior cingulate cortex and middle frontal sulcus, inferior and superior temporal areas, intraparietal sulcus, middle occipital sulcus and white matter (Fig.2b, Fig.3). This analysis revealed dissimilar spatial patterns with an overlap of a single channel between informed and arbitrary decision-making.

**Figure 3.**
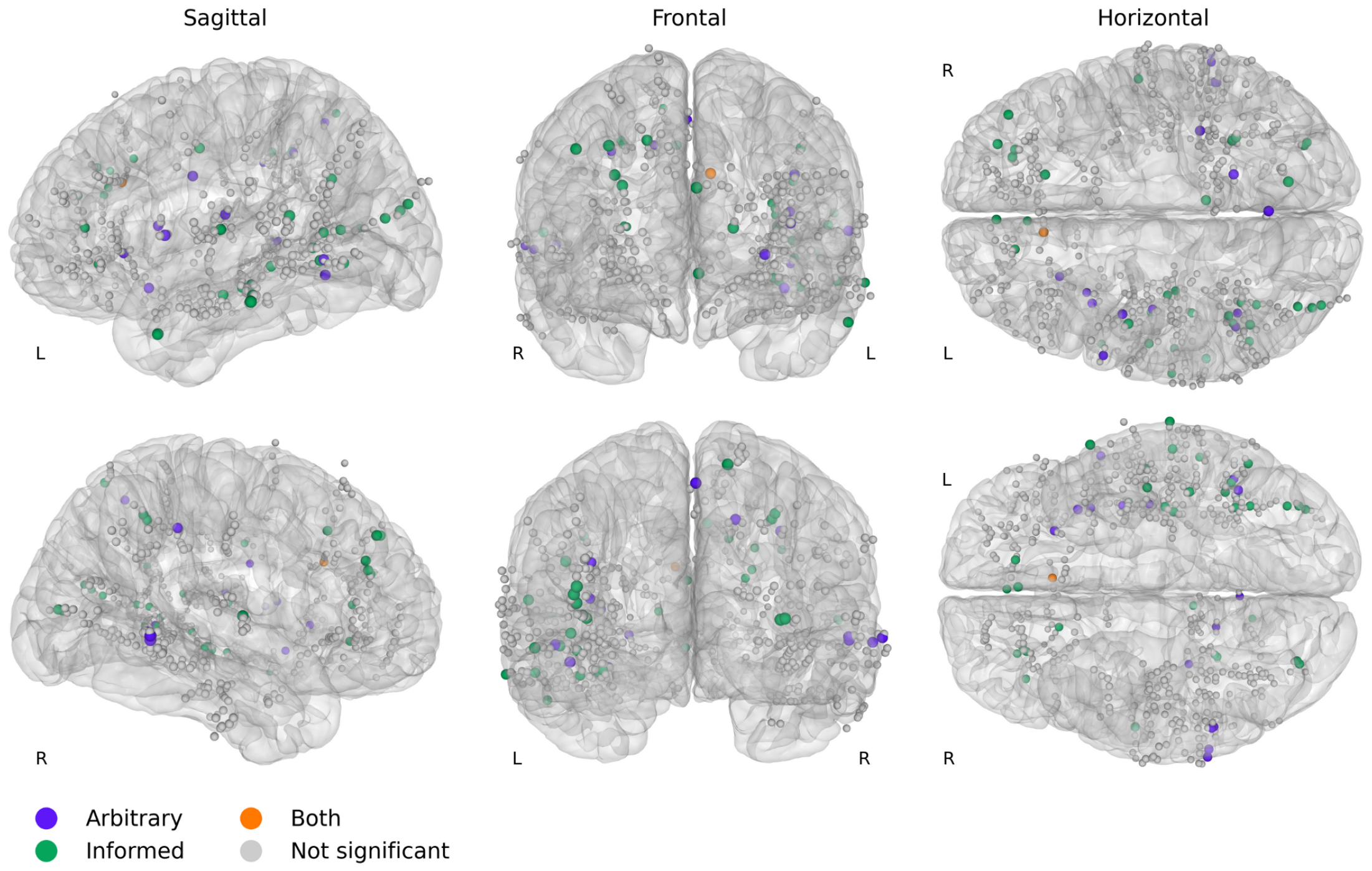
Spatial layout of informed and arbitrary decision decoding. Contacts anatomical location of all participants are shown in the MNI template space (obtained through warping from each individual’s native MRI space) from sagittal, horizontal and coronal views. Color indicates whether the channel resulted in significant decoding ROC AUCs (*p* < 0.05, permutation tests, T-Max correction) during arbitrary condition only (purple), informed condition only (green), informed and arbitrary conditions (orange) or none (grey).

### Time course of decision decoding

Lastly, we examined the temporal dynamics of decision decoding. To achieve this, we first split the 2s time window including inter-trial delay, stimulus, and cue periods, into 39 partially overlapping windows of 100ms. Then, for each window, an LDA classifier was applied to the average multi-band LFP power (Methods). Only the channels that had significant decoding performance in the previous analysis were included in this analysis. Permutation tests showed that a total of 31 (for informed decisions) and 15 (arbitrary decisions) channels reached significant decoding accuracy (ROC AUC, *p* < 0.05, permutation tests, T-Max correction) in at least 1 window. For further analysis, we selected the channels in which decoding accuracy was significant in at least 2 consecutive temporal windows during stimulus presentation. This criterion was met in 21 channels in classification of informed decisions and 7 channels in classification of arbitrary decisions.

As shown in Fig.4a and e, the time course of decoding accuracy varied substantially across channels. A linear regression analysis (Methods) revealed a significant interaction effect between time and condition on ROC AUC scores (*β* = 0.002, *SE* = 0.001, *t* = 2.89, *p* < 0.01), suggesting that the effect of time on accuracy differs across conditions, with overall higher scores in arbitrary compared to informed decisions at earlier time points.

**Figure 4.**
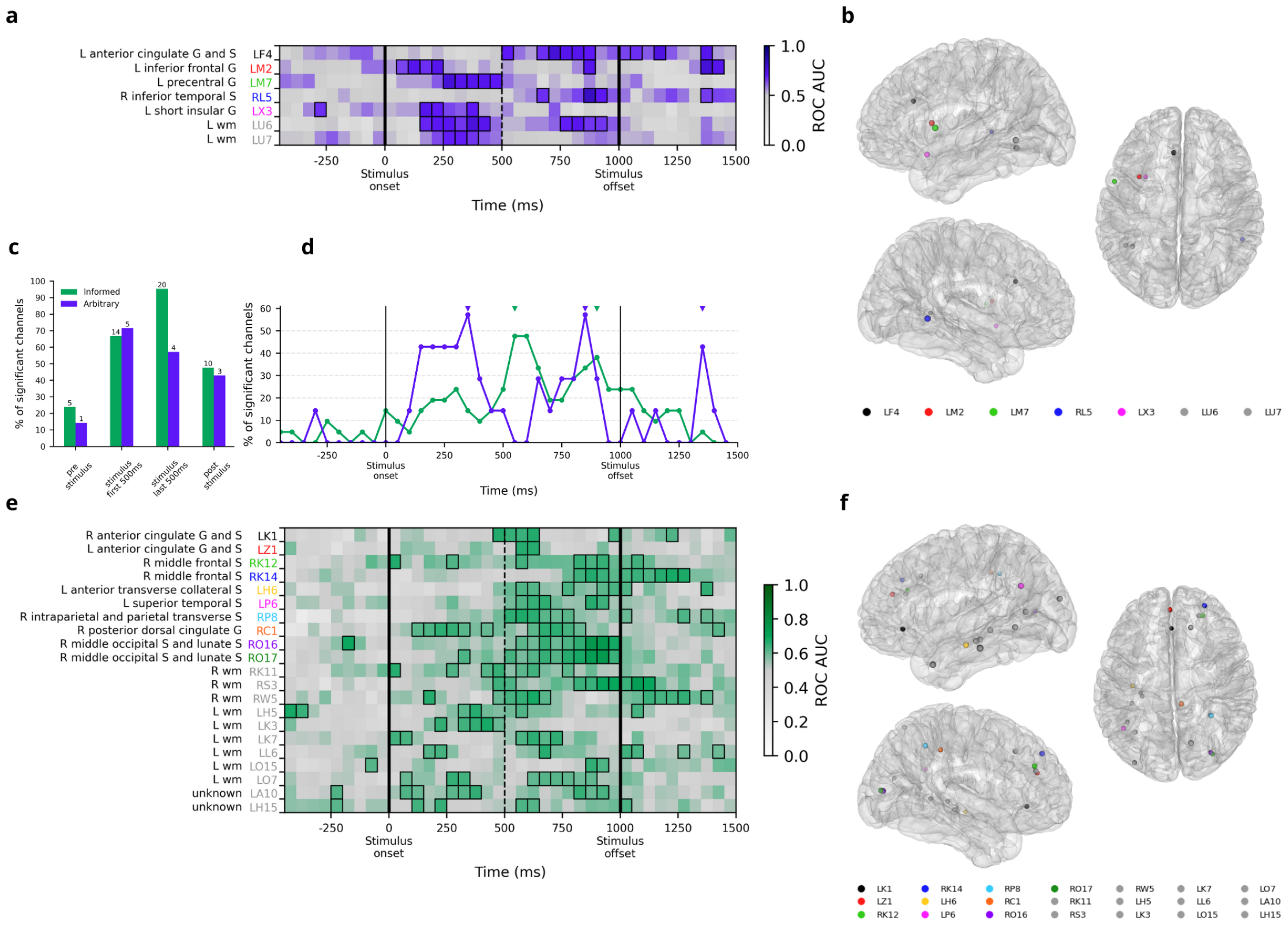
Spatio-temporal configuration of informed and arbitrary decision decoding. **a, *Time course of arbitrary decision decoding***. ROC AUC scores are plotted as a function of time (time-locked to stimulus onset) for each analyzed channel. Thick borders indicate the time windows in which ROC AUC is significantly higher than chance (*p* < 0.05, permutation tests, T-Max correction). Vertical lines signal stimulus onset and offset (full) and half-period of stimulus presentation (dashed). Colored channels’ labels represent channel anatomical location in **b**; grey labels indicate channels in white matter or unknown location. **b, *Spatial layout of arbitrary decision decoding***. Sagittal and horizontal view of the MNI template brain. Dots show the anatomical location of channels in **a** (color-coded). **c, *Percentage of significant channels across task periods***. Percentage and absolute number (above each bar) of channels showing significant ROC AUC (*p* < 0.05, permutation tests, T-Max correction) of informed (green) and arbitrary (purple) decisions for each task period. **d, *Percentage of significant channels across time***. Time-course of the percentage of channels showing significant ROC AUC (*p* < 0.05, permutation tests, T-Max correction) of informed (green) and arbitrary (purple) decisions. Triangles on top indicate peak locations for each condition. **e, f**, Same as in **a, b**, for informed decision condition.

To assess the time effect for each condition, we first examined the percentage of significant channels in each task period, for each condition. Most channels showed significant decoding accuracy during the last 500ms of stimulus presentation (*n* = 4 in arbitrary, *n* = 20 in informed) and within the first 500ms (*n* = 5 in arbitrary, *n* = 14 in informed). Additionally, in some channels we were able to decode upcoming decision during pre-stimulus fixation, preceding motor response by more than 1 second (*n* = 1 in arbitrary, *n* = 5 in informed). Finally, some channels showed significant decoding accuracy during the post-stimulus cue period (*n* = 3 arbitrary, *n* = 10 informed) (Fig.4c). These findings support the idea that LFP activity conveys information about upcoming categorization choice while the stimulus is visually perceived and close to the corresponding motor response, but also well before delivery of motor response.

Lastly, we focused on the percentage of significant channels at each time bin and identified the main peaks for each condition (Methods). We found three peaks for arbitrary decisions, located at 350ms, 850ms and 1350ms after stimulus onset, and two peaks in the informed condition, located at 550ms and 900ms after stimulus onset (Fig.4d). This analysis suggests that the time course of this representation differs between arbitrary and informed conditions, with predictive neural activity emerging earlier for arbitrary decisions compared to informed decisions.

## Discussion

We showed that individual category decisions can be decoded from single-channel multi-band LFP power, with a ROC AUC of up to 0.77. This means that, at the single channel level, the average power value of different LFP frequency bands - during stimulus presentation - provides valuable information to successfully predict whether the subject will choose the “Winning” or the “Losing” category. This suggests that broadband electrophysiological activity conveys information about choice, constituting a potential mechanism by which this information is represented in the brain.

Our task allowed us to distinguish two types of decisions. In one-third of the trials, participants had to make arbitrary choices on novel stimuli where the only possible strategy was to guess, by relying on reflexes or habits, as opposed to goal-directed instrumental learning. The remaining two-thirds of the trials involved previously seen stimuli, for which participants had to retrieve the correct category based on previous choices and feedback received, thus making informed decisions. In our paradigm, the sensory input and motor output remain constant across all conditions. Thus, distinctions between the arbitrary and informed conditions are solely attributable to variations in the available decision-making information provided to the agents. We explored decoding of single choices separately for each decision condition. Importantly, we found that both arbitrary and informed types of decisions could be successfully predicted, although their representation showed distinct spatial layouts, both in terms of the number of informative channels and in terms of brain areas involved.

First of all, arbitrary choices could be decoded from the activity of a relatively smaller set of channels (*n* = 16) compared to informed decisions (*n* = 35). Notably, the vast majority of these channels exhibited a specialized preference for the representation of a single decision condition. In fact, only one channel was informative in both conditions, while the remaining 49 channels selectively encoded either arbitrary or informed decisions.

Secondly, these two sets of specialized channels were located in distinct brain regions, allowing us to delineate spatial differences in the representation of each decision type. Channels encoding informed choices were distributed across a wide range of brain areas, including the anterior and posterior cingulate cortex, middle frontal gyrus, middle occipital gyrus, intraparietal sulcus, inferior temporal gyrus, superior temporal sulcus, transverse gyrus, and collateral sulcus. The second set, comprising channels representing arbitrary decisions, was located in the precentral gyrus, inferior frontal gyrus, inferior temporal sulcus, superolateral temporal gyrus, and insula. Lastly, the only one channel concurrently representing both decision types was located in the anterior cingulate cortex.

Hence, it appears that each decision type is distinctly encoded by a specific ensemble of widely distributed brain areas, while the anterior cingulate cortex stands as the sole region involved in the representation of both decision types. This is in line with previous studies that suggest that arbitrary and informed decision-making recruits distinct neural systems (***Bold et al., 2022***; ***Travers et al., 2021***; ***Maoz et al., 2019***; ***Travers and Haggard, 2021***).

Interestingly, we also found above-chance decoding of both types of decisions in white matter contacts, in agreement with previous results showing that white matter activity contains meaningful information for decoding tasks (***Mercier et al., 2017***).

In order to gain a deeper understanding of the temporal dynamics of decision representation, we examined the time course of decision decoding across the pre-stimulus fixation, stimulus presentation, and post-stimulus cue periods. Generally, the majority of channels exhibited the emergence of choice representation primarily during the final 500ms of stimulus presentation. Progressively fewer channels provided significant decoding in the initial 500ms of stimulus presentation, the post-stimulus cue, and the pre-stimulus fixation periods.

Importantly, we found that arbitrary and informed decisions exhibit different time courses of predictive power. While the number of channels with significant decoding accuracy first peaked at 550ms in informed decisions, for arbitrary decisions we identified an earlier first peak, located at 350ms. This observation aligns with a framework implying potential distinct mechanisms involved in the two processes (***Balleine, 2019***). Specifically, the early surge in predictive capability during arbitrary decisions may reflect the representation of reflexes or habits, whereas the later rise in informed decisions would reflect the recruitment of information from instrumental, goal-directed learning.

Both conditions showed a second peak located at 850ms (arbitrary) and 900ms (informed). Given the average timing of motor responses (335ms ±228ms following stimulus offset), it is conceivable that neural activity observed during the latter windows of stimulus presentation and poststimulus cue periods may reflect the preparation of motor acts. Potentially, channels exhibiting significant prediction accuracy during this timeframe may convey information related to motor planning rather than abstract stimulus categorization. Nevertheless, the motor acts employed in our paradigm shared the same effector (dominant hand) and were kinematically comparable, making it challenging to distinguish from sEEG activity.

Interestingly, some channels showed significant prediction accuracy during the pre-stimulus fixation period. The predictive value of neural information for responses, even before the stimulus was presented (Fig.4), is of particular interest, as it is likely associated with response biases and stimulus prediction. This could either be indicative of participants tracking the absolute occurrence of each stimulus, deploying statistical learning to detect potential underlying patterns of the loss-win distribution that would allow them to predict the next stimulus. Or it might be associated with a response bias.

Decision-making involves a complex interplay of multiple cognitive processes, including perception and memory. In our experimental setup, while informed and arbitrary decisions do not differ at the perceptual level, they do differ in the involvement of memory processes. Informed choices require the retrieval of previously stored information, whereas arbitrary choices may involve attempted but ultimately unsuccessful memory retrieval (due to lacking prior knowledge) and the encoding of new information (to establish stimulus-action mappings). The differences in the spatio-temporal patterns of decision representation that we report may stem from the differential engagement of memory processes in the two decision conditions (***Hong et al., 2023***; ***Rutishauser et al., 2021***). Future steps could focus on further exploring the role of memory processes in decision representation, by comparing memory encoding and retrieval within and outside a decisionmaking context.

In summary, our study demonstrated that the representation of categorization choice was observed prior to the motor response in both arbitrary and informed decisions. However, there were differences in the timing and spatial distribution of this representation between the two conditions, pointing out the involvement of distinct underlying mechanisms. These findings suggest that various aspects of decision-making, ranging from stimulus processing to action planning, are mediated by distinct mechanisms and brain regions.

To conclude, we show that both arbitrary and informed decision-making can be decoded from single-trial LFPs in sEEG recordings, which presents an important step towards neuroprostheses for decision processes and cognitive BCIs (***Wallis, 2018***). The brain-wide distributed recordings allow us to provide detailed information about the neural substrate of informed and arbitrary decision processes. Additionally, we bring robust evidence in support of the idea that single choices are represented by the brain through oscillations and shed further light on the spatial and temporal dynamics of arbitrary and informed decision-making.

## Methods and Materials

### Participants

Five patients (Tab.1) suffering from pharmaco-resistant epilepsy underwent implantation of sEEG electrodes for seizure source identification at the Maastricht University Medical Center. After surgery, the patients were transferred to the Epilepsy Center Kempenhaeghe to undergo the monitoring process. Here, patients agreed to voluntarily participate in our experiment. The study was approved by the Medical Ethics Review Committee of Maastricht University Medical Center and Epilepsy Center Kempenhaeghe and all patients provided written, informed consent for participating in the study.

**Table 1.** Sex, Age, and electrode contacts information for each participant. Note that the number of contacts shown here refers to the contacts from which sEEG activity was recorded and which were included in the analyses, not the total number of implanted contacts.

| Participant | Sex | Age | Shafts | Left Contacts | Right Contacts | Total contacts |
| --- | --- | --- | --- | --- | --- | --- |
| P01 | F | 52 | 12 | 61 | 66 | 127 |
| P02 | M | 36 | 13 | 0 | 122 | 122 |
| P03 | M | 30 | 12 | 98 | 59 | 157 |
| P04 | M | 36 | 15 | 109 | 18 | 127 |
| P05 | M | 21 | 6 | 57 | 0 | 57 |

### Electrode implants

Patients were implanted with platinum-iridium DIXI MicroDeep electrodes (DIXI Medical, France) with a diameter of 0.8 mm, a contact length of 2 mm, and an intercontact distance of 1.5 mm. Electrode placement was decided solely based on clinical needs of each patient. T1-weighted magnetic resonance imaging (MRI) scans were acquired pre-surgery for implantation planning. Electrode locations were confirmed post-surgery through co-registration of the pre-surgery MRI with a postoperative Computerized Tomography (CT) scan. A total of 127, 122, 157, 127, and 57 contacts were recorded for each participant (Tab.1, Fig.5). Anatomical labelling was conducted through cortical parcellation based on the Destrieux atlas (***Destrieux et al., 2010***) using img-pipe (***Hamilton et al., 2017***) and FreeSurfer (***Fischl, 2012***). The electrodes labelled as “Unknown” are usually located in areas that are not identifiable through FreeSurfer parcellation, such as electrodes positioned slightly outside the cortex, or at the border between gray and white matter. For visualization purposes only, contact locations in each individual’s native MRI space were warped to the average MNI space (***Mazziotta et al., 1995***). Note that for this reason, electrode positions in the MNI space might appear different from their respective labels.

**Figure 5.**
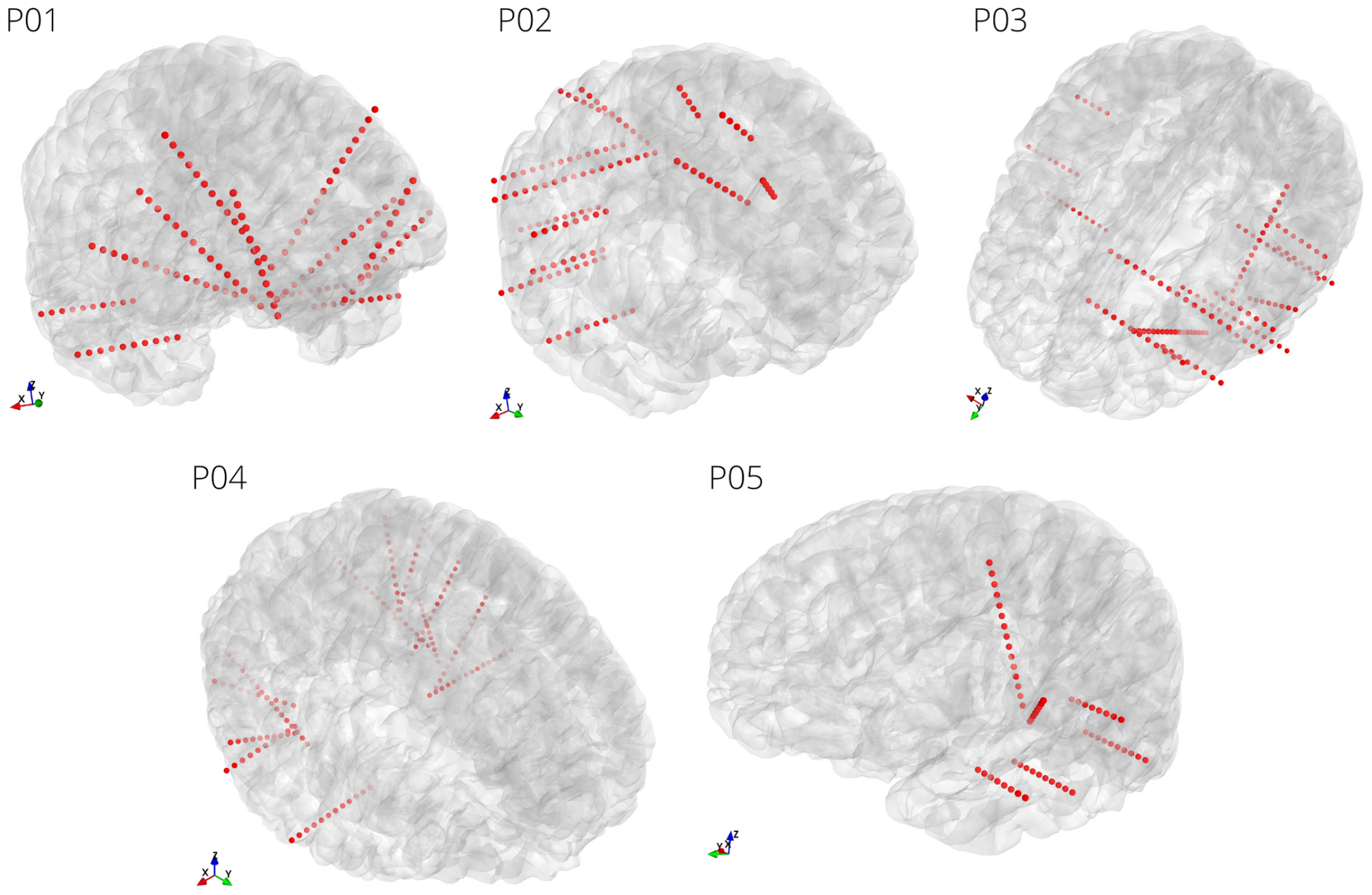
Electrode placement for all participants. For each participant, implanted electrodes are shown in the native MRI space. Electrode locations are determined through co-registration of pre-surgical MRI and post-surgical CT scans. Note that, based on clinical needs, data is recorded from only a subgroup of the implanted electrodes.

### Recording setup

Neural time series data were recorded with two or three stacked Micromed SD LTM amplifier (Micromed S.p.A., Italy) at 1024Hz sampling rate and referenced to a common white matter electrode. Neural data were synchronized to experimental events with LabStreamingLayer (***Kothe, 2014***).

### Task and stimuli

We designed a simple feedback-guided categorization task in which the goal is to assign each element of a set of visual stimuli to one of two abstract categories, adopting a trial-and-error strategy and minimizing error rates.

Stimuli were randomly chosen from the Novel Object and Unusual Name (NOUN) Database (***Horst and Hout, 2016***). The NOUN database contains images of non-real objects, complex and distinct entities that vary along several feature dimensions in addition to shape and color. This sort of stimulus allows the minimization of preconceived valence and intrinsic stimulus information. Stimuli were randomly assigned to one of two abstract categories, named “Winning”, referred to as “W”, and “Losing”, or “L”, in equal proportions.

Each experimental session consisted of 10 blocks of 9 trials, for a total of 90 trials per session. In each block, three stimuli were presented, each for three times, in a pseudo-randomized order (Fig.1).

Each trial began with a fixation cross presented centrally on a black computer screen. After 500ms, one stimulus was presented for 1000ms. Participants were asked to pay attention to the stimulus and to withhold any response until it disappeared. Then, a cue was presented for a maximum of 2000ms, during which participants were instructed to indicate the category associated with the stimulus, by pressing with their dominant hand the right (for category “L”) or left (for category “W”) arrow key on the keyboard to indicate the category they thought was associated with this stimulus. The cue disappeared when the button was pressed. Button presses before stimulus onset were not considered. If the participant did not choose in time, a warning to speed up was shown, and the trial was considered an error. At cue offset, the fixation cross reappeared for 500ms, followed by feedback presented for 1000ms. The feedback consisted of a colored screen, either blue or yellow, indicating whether the choice was correct or incorrect (i.e., whether the participants assigned the stimulus to its correct category). Additionally, the sound of coins dropping (for correct responses) or a buzzing sound (for incorrect ones) was presented. Right after the feedback presentation, the subsequent trial started. Each trial lasted between 3000ms and 5000ms, for a total experiment duration between 4.5 minutes and 7.5 minutes, depending on the participant’s speed. Stimulus-category associations were stochastic and unknown to the participant. Thus, participants were asked to first guess and then learn the correct categories based on the received feedback.

A training session with a different set of stimuli was performed for each participant to familiarize them with the task and ensure the instructions were clear and understood.

### Data preprocessing and features extraction

As a first step, sEEG signals were re-referenced using an electrode shaft referencing method (***Li et al., 2018***) where each contact is re-referenced to the mean signal of all contacts within the same electrode shaft. Then, data was filtered into 5 frequency bands: delta (1-3 Hz), theta (4-7 Hz), alpha (8-12 Hz), beta (13-30 Hz), high-gamma (70-120 Hz), by applying a zero-phase, 4th order Butterworth bandpass IIR filter. High-gamma was included as it is known to correlate with ensemble spiking (***Ray et al., 2008***) and provides localized information as shown in a number of intracranial EEG decoding studies (***Herff et al., 2020***). Subsequently, we estimated the envelope by extracting the magnitude of the analytic signal obtained through the Hilbert transform. Lastly, the obtained time series were epoched to extract time windows of interest for the subsequent analyses. First, a 1000ms window corresponding to stimulus presentation was extracted and used for the first decoding analysis. For the time course decoding analysis, the 2000ms epoch, including baseline (500ms, corresponding to fixation cross preceding stimulus presentation), stimulus (1000ms, corresponding to stimulus presentation) and response (500ms following stimulus offset) was segmented into 39 100ms windows with a 50ms frameshift.

### Behavioral statistical analysis

Accuracy was measured as the number of correct trials over total trials for each stimulus presentation. The goal of this analysis was to establish whether accuracy differed between decisions when stimuli were presented for the first time (arbitrary) and decisions in the second and third stimulus presentations (informed). To determine the effect of stimulus presentation on accuracy scores, we performed a one-way repeated measures ANOVA and Tukey’s honestly significant difference (HSD) test for multiple comparisons (*α* = 0.05). Reaction times were analyzed considering the first arrow press after stimulus onset, therefore including also early responses (arrow press before cue onset). The number of trials with early responses was variable across participants (7, 50, 8, 5, 28 out of 90 trials, respectively, for P01, P02, P03, P04, and P05), occurring on average 250ms ±93ms before stimulus offset.

### Decoding analysis

The aim of this analysis was to explore whether we could predict upcoming decision (“W” or right presses vs “L” or left presses) based on multi-band LFP power of each single channel. We performed this analysis separately for the arbitrary decision condition (first stimulus presentation, 30 trials) and for the informed decision condition (second and third stimulus presentation, 60 trials). For each channel, we run a decoder using Linear discriminant analysis (LDA) with shrinkage. The feature space was obtained by averaging the power of each of 5 frequency bands over the entire stimulus window (1000ms), resulting in a matrix of dimensionality = trials (30 or 60, depending on the decision condition) by frequency bands (5). Subsequent response (“W” or “L”) for each trial constituted the target variable. Each decoder was evaluated with the area under the Receiver Operating Characteristic curve (ROC-AUC) using leave-one-out cross-validation.

This decoding analysis allowed us to determine which channels exhibited activity patterns containing information about the decision.

In a second step, we aimed to explore more in-depth the time course of decision decoding in these channels. We repeated the previous analyses for each of the 39 time windows obtained by the segmentation of the 2000ms period including baseline, stimulus presentation and response (see Data preprocessing and feature extraction), for arbitrary and informed decisions separately. Here, features consisted of the average power over the 100ms windows, for each frequency band and trial. All decoding analyses were implemented in Python with scikit-learn package (***Pedregosa et al., 2011***).

### Permutation tests

To assess the statistical significance of our decoding performance, we employed non-parametric permutation tests (***Ojala and Garriga, 2010***). For each participant and decoding analysis, we first estimated the null distribution under the assumption of no dependency between the extracted features and the target variable. For the first decoding analysis (average over the entire stimulus period), this was achieved by estimating the decoding ROC AUC with leave-one-out cross-validation on permuted labels 1000 times for each channel. This process yielded a distribution of 1000 randomized sample decoding results for each channel, effectively capturing the variability under the null hypothesis. On the other hand, for the time-course decoding analyses, the null distribution for each channel was estimated by randomly selecting one among the 39 time-windows, permuting the labels and performing the decoding analysis for a total of 1000 times.

Subsequently, we first computed an individual significance threshold for each channel by identifying the 95th percentile of its respective null distribution. Then, to correct for multiple comparisons, we determined the maximum threshold across all channels. This maximum threshold served as our final significance threshold for all channels, ensuring a stringent control of the familywise error rate. Only decoders that got an observed ROC AUC value higher than the maximum threshold were considered significant.

### Additional analyses

To test for differences in decoding accuracy between conditions over time, we conducted a regression analysis, modeling ROC AUC as a function of Time and Condition (*R*^2^ = 0.051, *F* (3, 1088) = 19.43, *p* < 0.001). The results revealed a significant main effect of condition on ROC AUC scores (*β* = −0.04, *SE* = 0.01, *t* = −3.13, *p* < 0.01), a non-significant main effect of time on accuracy (*β* = 0.0005, *SE* = 0.000, *t* = 0.97, *p* = 0.33), and a significant interaction effect between time and condition on accuracy (*β* = 0.002, *SE* = 0.001, *t* = 2.89, *p* < 0.01).

In order to further depict the specific differences in the time course of decoding accuracy between conditions, we conducted peak detection analysis on the percentage of significant channels over time bins. We defined peaks as the local maxima with a minimum height of 35, for each condition.

## Acknowledgments

This publication is part of the project DESIS (with project number VI.Veni.194.021) which was awarded by the Dutch Research Council (NWO) to C.H.

## Additional information

### Data Archival

Code used in this study is available at https://github.com/LauraMarras/DecodingDecisionsSEEG. All data used in this study is available on https://osf.io/uaq4n/.

### Author contribution

L.M., M.L.F.J., S.A.H and C.H. designed the experiment. M.V., J.P.D., and M.C.O. recorded the data. S.T., L.W., A.J.C, and P.K. included the participants. M.V. and S.G. localized the electrodes. L.M. and C.H. analyzed the data and wrote the manuscript. All authors reviewed the manuscript.

